# Haisu: Hierarchical Supervised Nonlinear Dimensionality Reduction

**DOI:** 10.1101/2020.10.05.324798

**Authors:** Kevin C. VanHorn, Murat Can Çobanoğlu

## Abstract

We propose a novel strategy for incorporating hierarchical supervised label information into nonlinear dimensionality reduction techniques. Specifically, we extend t-SNE, UMAP, and PHATE to include known or predicted class labels and demonstrate the efficacy of our approach on multiple single-cell RNA sequencing datasets. Our approach, “Haisu,” is applicable across domains and methods of nonlinear dimensionality reduction. In general, the mathematical effect of Haisu can be summarized as a variable perturbation of the high dimensional space in which the original data is observed. We thereby preserve the core characteristics of the visualization method and only change the manifold to respect known or assumed class labels when provided. Our strategy is designed to aid in the discovery and understanding of underlying patterns in a dataset that is heavily influenced by parent-child relationships. We show that using our approach can also help in semi-supervised settings where labels are known for only some datapoints (for instance when only a fraction of the cells is labeled). In summary, Haisu extends existing popular visualization methods to enable a user to incorporate known, relevant relationships via a user-defined hierarchical distancing factor.

**Availability:** github.com/Cobanoglu-Lab/Haisu

## Introduction

Dimensionality reduction (DR) involves the elimination of features or random variables of a given dataset. DR is a crucial component in modern analysis techniques and is widely applicable due to the rapid growth in data available to businesses, scientists, and public administration [1]. This data can be comprised of text-based or multimedia content from entertainment, research, and business sectors. The role of DR for feature selection and extraction and data preprocessing is especially pertinent when analyzing immense volumes of data with methods such as machine learning [2].

We focus on the use of dimensionality reduction for visualization of biomedical data. When high-dimensional data is difficult to summarize, DR is often effective for transforming the data to two or three dimensions. This can be achieved through linear and nonlinear approaches. Nonlinear dimensionality reduction (NLDR) is preferable for capturing the local and global structure of the data, where linear DR tends to be faster and more effective solely for global patterns [2]. Our method, Haisu, introduces a general-purpose NLDR extension based on three popular visualization methods that incorporates an input hierarchy. Haisu can be manually weighted with a strength factor to better represent patterns in the data at any degree of influence. We implement this technique for both known and predicted class-based hierarchies with the option to influence the strength of a predicted class’ structural effect based on its likelihood.

Many nonlinear techniques exist for dimensionality reduction and are mainly unsupervised. Examples of such methods include Sammon mapping [3], Curvilinear Components Analysis (CCA) [4], Stochastic Neighbor Embedding (SNE) [5], Isomap [6], Maximum Variance Unfolding (MVU) [7], Locally Linear Embedding (LLE) [8], and Laplacian Eigenmaps [9]. These are often effective with artificial data but struggle to maintain both local and global structure on real-world data [10]. For this reason, we primarily target modern nonlinear dimensionality reduction methods. For demonstrational purposes, we show the effect of Haisu on Principal Component Analysis (PCA) as well, a widely used unsupervised linear technique.

To our knowledge, no modification has been introduced to enable a general-purpose NLDR technique to be influenced by a variable-strength hierarchy. Auxiliary kernel-based semi-supervised techniques can be effective at preserving the local and global structure of data and have recently shown effective in sc-RNA sequencing data [11]. Haisu is independent of such pre-processing method and can even be used in a complementary manner, preserving their effects while introducing a supervised hierarchy-based component. Other supervised and semi-supervised DR approaches exist but are not readily applicable to general-purpose visualization [12,13]. Finally, separate techniques that integrate hierarchical information such as Hierarchical Manifold Learning are less concerned with visual analysis in lower dimensions and thus not as relevant to our work [14]. We aim to provide a context-independent approach that incorporates a tunable supervised hierarchical influence across stochastic embedding approaches. We argue that if a known hierarchy exists for a dataset, it can provide benefit when integrated into the analysis. For example, such an approach can help to alleviate the case where important and explicit relationships in the data are ignored due to low expression.

To explore the benefits of integrating a supervised hierarchical approach for DR, we primarily focus on single-cell RNA (sc-RNA) sequencing datasets in this manuscript. Nonlinear dimensionality reduction is widely applied to sc-RNA sequencing, and many such datasets are influenced by an interpretable cell-based hierarchy known prior to experimentation. Because clusters identified via NLDR are frequently integrated into unsupervised cell-type prediction models, we suggest that Haisu is particularly useful in this domain [15–17]. We highlight three state-of-the-art NLDR approaches: t-SNE, UMAP, and PHATE.

Formulated by Maaten and Hinton, t-SNE is currently the premier dimensionality reduction method used for high-dimensional data visualization [18]. This technique is a variation of Stochastic Neighbor Embedding, or SNE, which converts the Euclidian distance between high-dimensional data points into conditional probabilities that represent similarities. T-SNE’s variation adds a Student’s t-distribution to SNE which solves the crowding problem that standard SNE suffers from. Uniform Manifold Approximation and Projection (UMAP) is competitive to t-SNE and aims to better preserve the global structure of data with better scalability. Finally, potential of heat diffusion for affinity-based transition embedding (PHATE) is a domain-specific visualization strategy designed for biological datasets [19]. PHATE aims to improve denoising and provide biological insight across datasets and data accession methods.

## Methods

### The Effect of Haisu on t-SNE, UMAP, PHATE

Given a high dimensional input 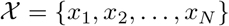, the asymmetric similarity ***s***_*j|i*_ between *x_i_, x_j_* is commonly influenced by a variable distance function *d*(*x_i_, x_j_*). Our algorithm introduces an additional pairwise distance modification *θ_ij_* based on the input hierarchy where 0 ≤ *θ_ij_* < 1. At lower values of *θ_ij_*, the distance *d*(*x_i_, x_j_*) is preserved, and at higher values, distances are penalized based on the corresponding hierarchy graph. We describe the repercussions of this modification to t-SNE, UMAP, and PHATE where ***s***_*i|i*_ = 0.

T-SNE extends the pairwise similarity definition ***s***_*j|i*_ from SNE to a conditional probability *p_j|i_* where 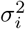 is the variance of a Gaussian centered at *x_i_*. T-SNE then determines joint probabilities *p_ij_* in high-dimensional space as symmetrized conditional probabilities s.t. *p_ij_* = *p_ji_*. With Haisu, the proportionality of *θ_ij_* to ***s***_*j|i*_ is inversely exponential because *θ* is a fractional modifier bound between 0 and 1. Thus as *θ* decreases with closer relative distances in the hierarchy graph, *d*(*x_i_, x_j_*) will decrease in the kernel, resulting in a higher modified similarity 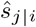. With the introduction of the hierarchical distance factor *θ_ij_*, we also include the resulting conditional and joint probabilities 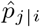 and 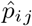 respectively:

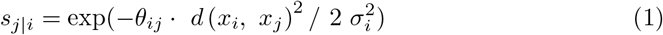

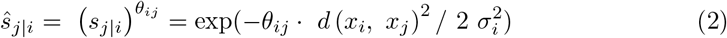

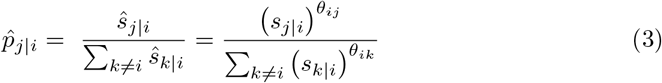

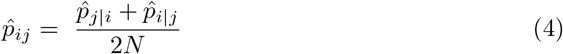

Pairwise similarities in UMAP are calculated as smoothed nearest neighbor distances where *n_i_* is the distance to the nearest neighbor of *i*, and 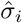 is the Haisu-modified normalizing factor. Without further normalization, UMAP then symmetrizes ***s***_*j|i*_ with a probabilistic fuzzy set union to obtain *p_ij_*.

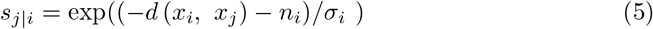

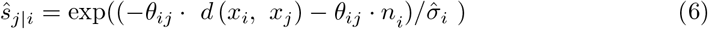

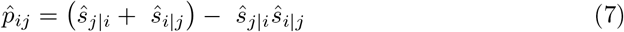

To compute the standard normalizing factor *σ_i_*, UMAP performs a binary search for *σ_i_* s.t. 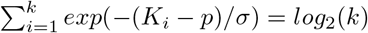. This optimization to *log*_2_(*k*) is an empirical measurement by UMAP to smooth the k-nearest distances *K*. Supplying a precomputed distance matrix modified by Haisu affects the calculation of the normalizing factor *σ_i_* via the distance to the nearest neighbors *K* which affects *p* = *K*_1_. Thus, via 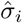, UMAP takes the hierarchical matrix *θ* into account when smoothing the *k* nearest neighbor distances for the pairwise similarity 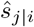. This is necessary to fix the cardinality of the fuzzy set of 1-simplices to a fixed value influenced by *θ*. Here the matrix *θ* is defined by *θ_ij_* across high-dimensional vectors from 1 to *N* assuming *x_i_, x_j_* ∈ *X* can be assigned labels (i.e. they are samples not features).

PHATE uses an α-decaying kernel function with an adaptive bandwidth *σ_i_* (for a gaussian kernel, *σ_i_* = 2). To better preserve global relationships, PHATE computes a diffusion geometry. We can apply *θ_ij_* to the distance function *d*(*x_i_, x_j_*) as with t-SNE and UMAP.

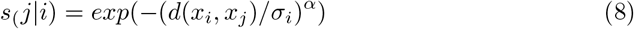

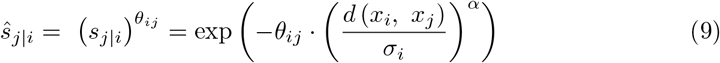

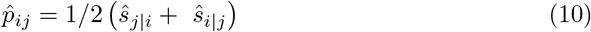

PHATE integrates the graph-tools library [20] where *α* is applied to the precomputed distance matrix, thus, the default effect of Haisu on ***s***_*j|i*_ is as follows which more closely preserves the effect of the decay factor *α*. As a byproduct, this results in a less drastic hierarchical distancing effect for higher values of *α*.

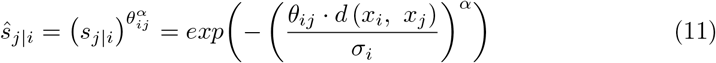

In review, the pairwise similarities for each NLDR method are formulated in a manner that the graph-based pairwise factor *θ_ij_* does not compromise a method’s primary architecture. Haisu modifies the transformation from global distances to local similarities solely in the pairwise distance function *d* (*x_i_, x_j_*). Because our method is subtractive, the structure of a given method’s lower dimensional embedding is identical to its unadulterated configuration when our strength factor *str* = 0. Finally, Haisu does not interfere with pairwise similarities in the low-dimensional embedding when high dimensional distances are not referenced. This further ensures that the appearance and structure of a Haisu embedding closely match those of its respective NLDR approach at any dimension. We will now describe the calculation of our hierarchical distancing factor *θ_ij_*.

### Hierarchical Supervised NLDR

Given an undirected (not necessarily connected) graph *G* with labels assigned to each node, we process input points *x_i_, x_j_* in high-dimensional space with labels *i*′, *j*′ respectively. We then find the shortest path distance of two undirected nodes 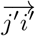. If the nodes are in different connected components, we then return zero as the path length. Define *maxdist* as the maximum length shortest path in *G*, in other words *maxdist* is defined as *max*(*S*) where *S* is the set of possible shortest paths in *G*. Finally, let *str* denote the strength of this function’s effect on the pairwise-distance function *d*(*x_i_, x_j_*). Given that *str* ∈ [0,1) the function is then bounded by [*str*, 1). The resulting function is as follows:

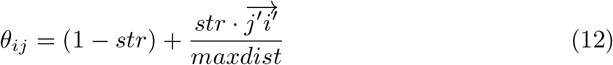

Using this formulation, a user can choose to input the adjacency matrix of the input graph *G* and the strength of the hierarchical distancing factor. Graphs can be specified for either dimension of the input dataset. In the case that there is a known hierarchy for the dataset, but labels are unknown, we specify a feature-based method that weights the strength of a predicted class’ structural impact based on its likelihood. Given the probabilities *p_i_* and *p_j_* of labels *i*′ and *j*′ for high-dimensional points *x_i_, x_j_*, we define *m* = *min*(*p_i_, p_j_*). In practice, this choice of probability could be *max, mean*, or any method of aggregation. We then weight the distancing factor *θ* with *m*:

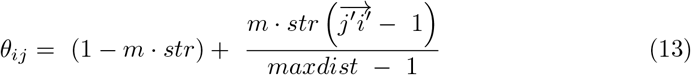

Our reference software is implemented in Python 3. We provide a modified pairwise distance matrix to any NLDR approach that accepts a ‘precomputed’ metric. The input parameters for our implementation include an adjacency list defining the class hierarchy, the strength factor str, a standard pairwise distance function from the scikit-learn library [21], a list of class labels, and a list of class probabilities when relevant.

We illustrate the effect of Haisu on PHATE, t-SNE, and UMAP at multiple strength factors, and include PCA to express more simply how Haisu modifies the pairwise distance matrix. Strength values of 1 are included for demonstrational purposes but are not usually informative due the inverse exponential nature of Haisu’s distancing factor *θ*. In the case of PHATE, this is especially evident due to the exponential effect of the decay factor *α* on *θ_ij_*. Thus, for the figures presented, we pass 0.999 as the strength value instead of 1 where indicated*.

## Results

We analyzed the performance of our technique on single-cell RNA-sequencing datasets of three different modalities. First, we include fresh peripheral blood mononuclear cells (PBMCs) from Zheng et al. to differentiate between T Cell subtypes based on function [22]. Second, we apply HAISU to a proximity-based hierarchy for embryonic cardiac cells [23]. Last, we include a hierarchy based on epithelial differentiation in healthy and ulcerative colitis patients [24]. Finally, to demonstrate the efficacy of our approach, we progressively remove TA 1 cell labels from the epithelial dataset. We denote str as the strength of Haisu’s hierarchical distancing factor *θ* as defined in the methods section.

We chose to target the high-dimensional single-cell dataset specified in Zheng et al. due to the ambiguity between T cell subclasses in the raw t-SNE plot. CD4+ and CD8+ T cells in [22] are highly interspersed making it difficult to discern relationships between the subclasses. Additionally, there is a known relationship between the given classes [25], which motivated the application of a hierarchy-based model. Our visualizations are produced from the filtered “Fresh 68k PBMCs (Donor A)” dataset provided by 10x Genomics [26]. Prior to analysis, UMIs (unique molecular identifiers) under a target variance were removed (we chose 0.1) and then min-max normalized by gene. We then randomly subsampled to 30,000 cells. The graph that we specify for this dataset is derived from a gene-based hierarchy with explicit classes from Zheng et al and implicit parent classes such as “Lymphoid” and “Conventional T Cell” (Fig. 1). We uphold parent-child relationships in the graph based on lineage and a dense set of nodes to distinguish between T cell types based on function. Through a purely lineage-based method, CD34+ should be in a different connected component, but we chose to keep it linked to the head node of the tree due to its early role in hematopoiesis.

**Figure 1.**
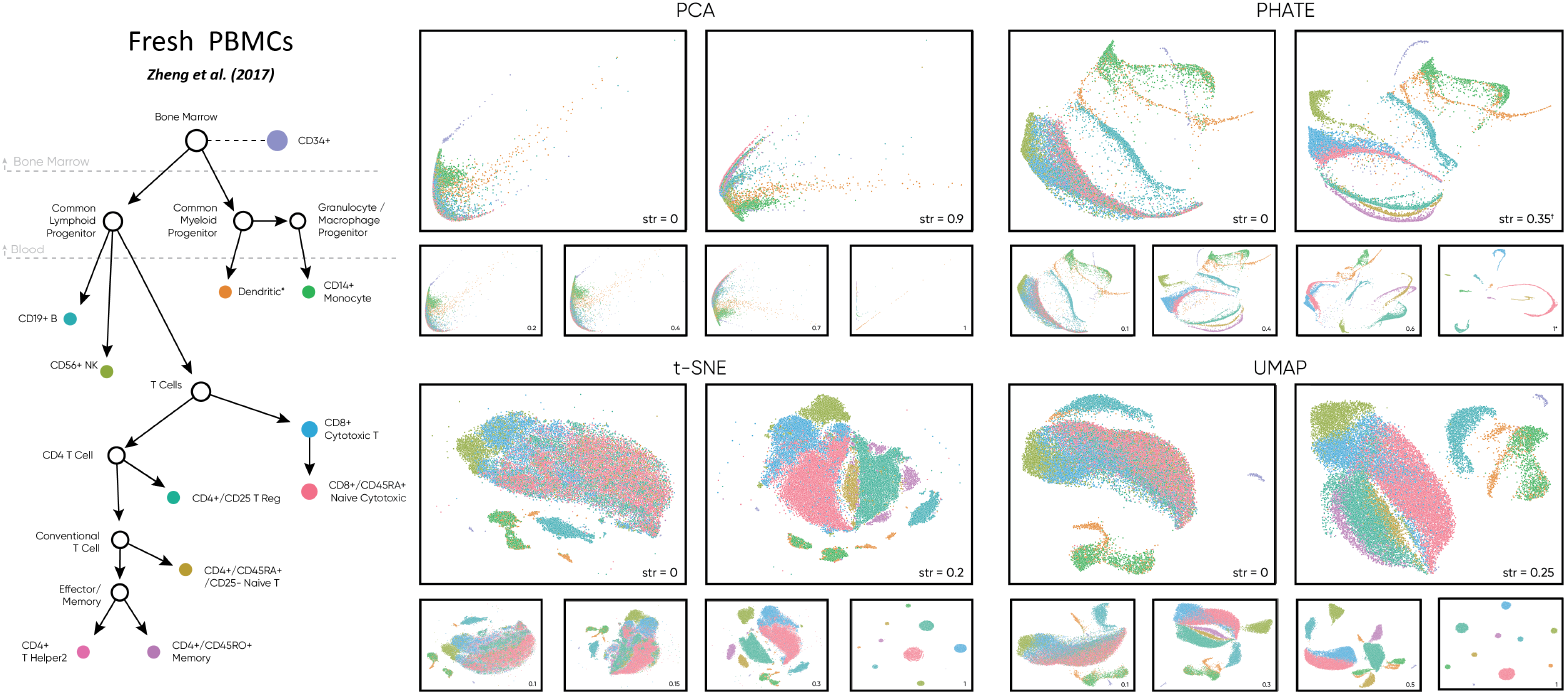
For this dataset, Haisu applies an input hierarchy based on cell function and lineage to guide the identification of sub-clusters. We display the effect of our method on popular nonlinear DR approaches and PCA at multiple ‘strength’ values (*str*), a tunable factor between 0 and 1 to control the strength of our hierarchical distancing function. Compared to raw NLDR (*str* = 0), Haisu reveals sub-clusters of T cells and better expresses the subtle relationship between datapoints in each method. †: Reflected on x-axis for visualization.

Across NLDR methods, Haisu effectively separates different populations of T-cells. For this PBMC dataset, lower strength values are more effective due to the lower inter-class variability of the processed UMI counts within the presence of outliers that decrease the size of the primary area of interest. By modulating the strength factor, we observe multiple effects on the data across DR methods. Firstly, original clusters of Monocyte, B, and Dendritic cells maintain relative distance and shape. Even when clusters of Monocytes migrate, local dendritic embedding remains consistent for strength values roughly less than 0.5. We also observe that Cytotoxic T, NK, and Naïve Cytotoxic T cells remain closely intermingled until higher strength values where NK cells cluster nearby due to the hierarchical distancing factor. Finally, a group of Cytotoxic T cells consistently groups with neighboring Naïve Cytotoxic cells in PHATE (*str* = 0.35), UMAP (*str* = 0.25), and t-SNE (*str* = 0.2, 0.3). This suggests that some Cytotoxic T cells are more like those labeled “Naïve Cytotoxic” than their own class and thus possibly mislabeled. Haisu in this effect can be used to encourage the re-evaluation of the prediction model for a given class. The Naïve T cell population which is difficult to observe in raw t-SNE and UMAP clearly borders Naïve Cytotoxic and T Regulatory cells at *str* = 0.2 and *str* = 0.25 respectively. This global relationship breaks down at higher strength values due to the stronger influence of sibling nodes (see UMAP at *str* = 0.3 and t-SNE at *str* = 0.3), highlighting the importance of choosing a correct lower-influential strength value to discern subtle relationships with Haisu. Lastly, our hierarchical t-SNE (H-tSNE) implies two or more subpopulations of Monocytes and CD4+ Memory T cells at *str* = 0.2.

We include the embryonic cardiac cell dataset for its inclusion of a proximity-based hierarchy that includes labels that correspond to the tissue region from which a cell sample was extracted [23]. Thus, the hierarchy is more tangible/concrete than one where classes are determined by gene expression such as in (Fig. 1). Because we do not include the unsupervised classes but rather the anatomical site hierarchy from Li et al, the graph-label assignment is truer to this mode of Haisu where label likelihood is not given. We suggest that our method is particularly valuable in this case where a gene-based embedding can be influenced by the relative distance between cell extraction sites. We include all samples with the tissue tag listed in the graph, drop genes under a variance of 0.5, and perform min-max normalization per cell.

For this dataset, there are two primary observable clusters in each raw NLDR method. This separation is maintained at lower strength values in each method where *str* ≤ 0.3. By introducing the hierarchical distancing factor, the ventricle subgroup is more discernable in both clusters. With NLDR techniques that maintain global structure more effectively such as PHATE and UMAP, Haisu is more informative between hierarchically distanced clusters. PHATE at *str* = 0.5 branches into three groups with two sub-groups for each same-label cluster. In this manner, Haisu rearranges the original graph based on the hierarchy but maintains the original global structure. For example, the atrioventricular canal cells are in each of the two primary clusters for the raw graph and in PHATE at *str* = 0.5, they are in a branch but maintain relative distance. UMAP at *str* = 0.4 demonstrates a similar effect but without the branching common to PHATE. For t-SNE, cells with siblings in the graph are too dramatically separated, and distances between the resulting clusters are less informative since t-SNE intra-cluster distances is not significant.

**Figure 2.**
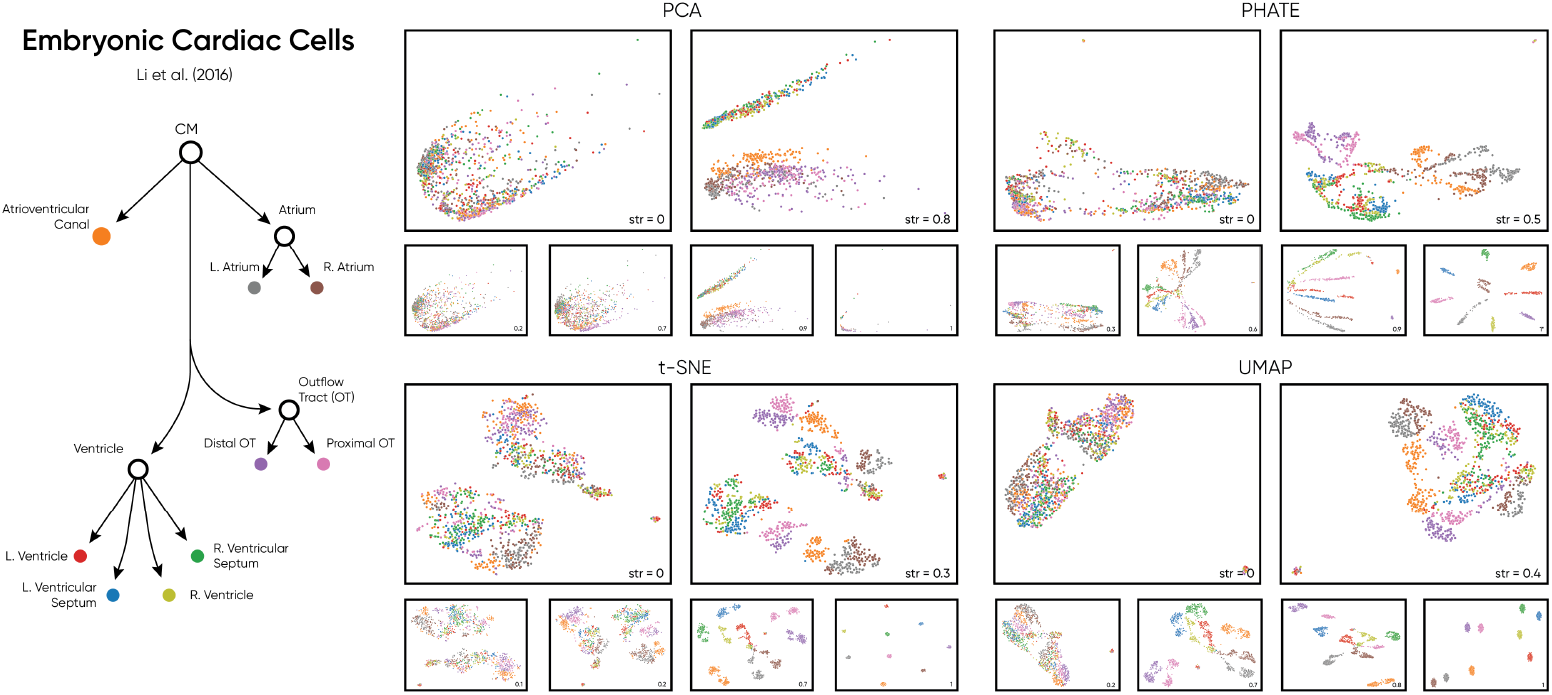
Haisu applied to anatomical embryonic cardiac cell subpopulations via a proximity-based hierarchy. The raw embeddings of each method indicate two primary clusters with cell label assignments that are spaced out within each cluster. Haisu helps to add clarity to the embedding in a manner true to the known external hierarchy. Labels are assumed to be 100% accurate as they are location-based, but anatomic regions can have similar transcriptomic profiles. Thus, Haisu in this context, factors in gene expression and location when determining a lower dimensional embedding at an appropriate strength.

**Figure 3.**
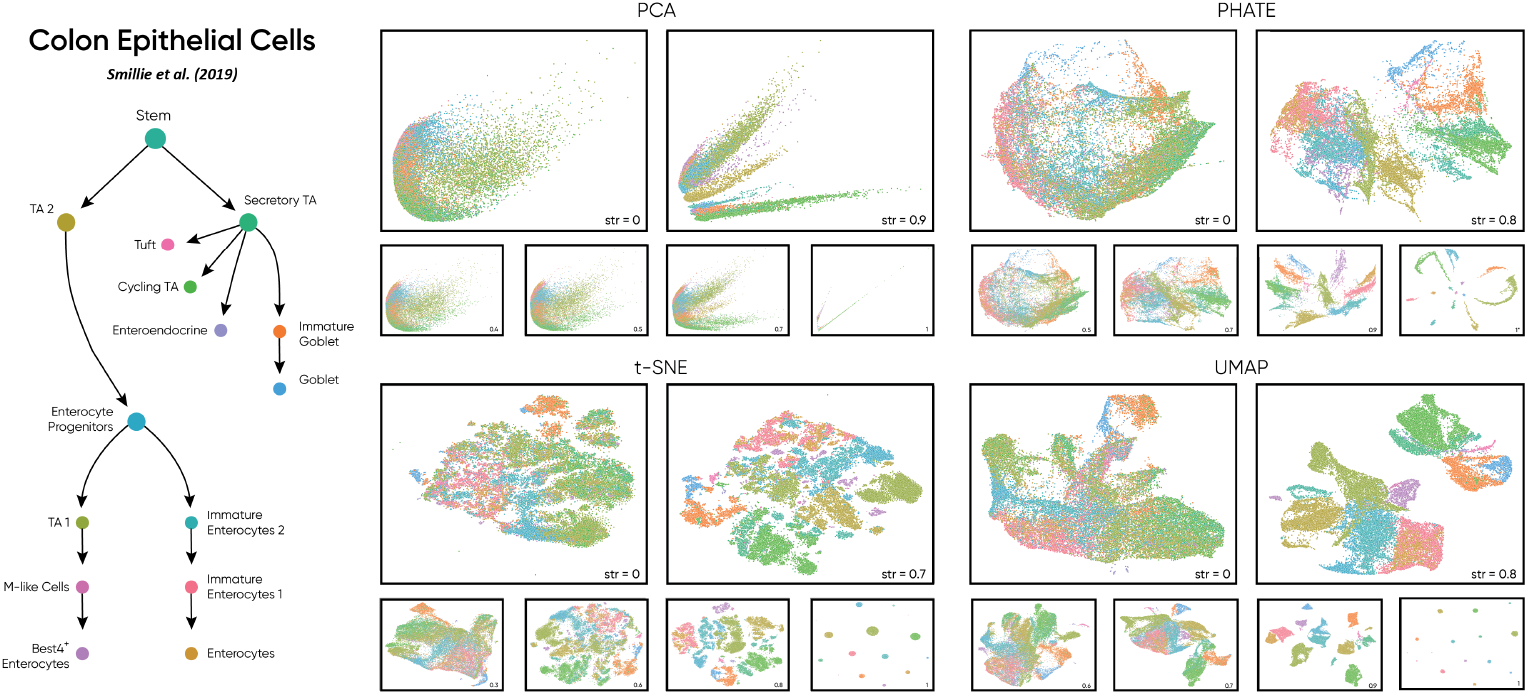
We illustrate the effect of Haisu within the context of an epithelial differenti-ation hierarchy in the context of healthy and ulcerative colitis patients. In this dataset, strength factors up to 0.8 uphold appearance of the raw embedding. Thus, with sufficient confidence in cell type labels, Haisu preserves the structure of the NLDR method while also allowing a simpler examination of more subtle inter-cluster relationships.

Finally, we investigate the application of Haisu to an epithelial differentiation dataset for cells during ulcerative colitis (randomly subsampled to 30,000 cells) [24]. We filtered out genes under a variance of 1 and performed min-max normalization across cells on the resulting matrix prior to subsampling. Without our method, the raw NLDR embeddings are too heterogeneous to discern any patterns in the data. Thus, we integrate the known hierarchy to create a more readable embedding. Most notably, by applying the hierarchy, we can observe sub-clusters that are not identifiable in the raw embeddings. Examining PHATE and UMAP at str=0.8, we observe two primary groups of cells caused by two primary parent nodes in the graph: secretory TA cells and enterocyte progenitors. Just as with the cardiac dataset, we expect DR techniques that preserve global structure to better reflect properties of the input graph. Prior to the clumping of graph node siblings at *str* = 0.9, PHATE and UMAP allow for the inspection of more subtle inter-cluster relationships such as the position of Best4+ Enterocytes among TA 1, M-like cells, enterocyte progenitors, and immature enterocytes 2 in PHATE *str* = 0.8. This suggests possible subpopulations or possible alternatives for the labeling of specific Best4+ Enterocytes that gravitate more toward the different mentioned classes. Smaller populations of cells such as M-like cells are also easier to view as the strength factor is increased. UMAP at *str* = 0.8 reveals a group of TA 1 cells that are placed within the large group of immature enterocytes 2, this relationship is too loose to observe in the raw UMAP embedding at str = 0. We similarly observe this same relationship in t-SNE at Str = 0.7 as immature enterocytes are spread among smaller TA 1 clusters. Though t-SNE is less effective at global relationships, it can be employed in this manner, finding cluster of cells with more than one class of cell, suggesting that the two (or more) share similar traits.

To empirically test the efficacy of Haisu’s ability to preserve the structure and relationships between the modified embedding and the original, we illustrate the effect of removing a class label. In Fig. 4, We progressively replace the label of TA 1 cells with a ‘dummy’ label, such that cells with the original label “TA 1” are completely removed from the influence of our hierarchical distancing factor. 100% removal indicates that all TA 1 cells in high-dimensional space are given the ‘dummy’ label. We suggest that even in an embedding highly influenced by the input graph (i.e., a higher value of str) Haisu maintains the relative positioning of datapoints even when they are unlabeled. This is a critical observation to communicate the efficacy of our approach such that Haisu at low strength values is a non-destructive method to influence an embedding based on a hierarchy.

In Fig. 4. TA 1 cells are colored a darker gray when their label is provided to HAISU, and a lighter gray when it is disclosed and given the label ‘dummy.’ Observing the relative position of TA 1 cells in the raw graphs of each NLDR method, we focus on a strong coupling with TA 2 Cycling TA cells closely distanced from the enterocyte progenitor cells. The TA 2 cell label is the grandparent of TA 1, so local clustering in t-SNE and global closeness in PHATE and UMAP is expected. This same relationship is expected to hold with enterocyte progenitors with are rewarded in Haisu because their cell label is the parent of ‘TA 1.’ However, Haisu does not encourage close positioning of TA 1 cells with Cycling TA cells in the embedding due to their distance in the graph. Thus, it is a remarkable testament to the preservation of global structure when we observe clustering of unlabeled TA 1 cells alongside Cycling TA. This is especially apparent in PHATE at *str* = 0.9, t-SNE at 1, and UMAP at 0.9 when 100% of TA 1 labels are removed. In PHATE at *str* = 0.9, we highlight the difference between 0% removal and 100% removal. In the former, Cycling TA cells are distanced into their own cluster, and TA 2/enterocyte progenitors remain close due to their node distance in the hierarchy. However, in the latter at 100% removal, we see that Cycling TA has returned to the same relative position as in the raw PHATE graph, next to TA 1. We outline this with a dotted shape in Fig. 4. At 100% removal we also observed that the unlabeled TA 1 cells remain close in the embedding to enterocyte progenitors across strength values and NLDR techniques.

**Figure 4.**
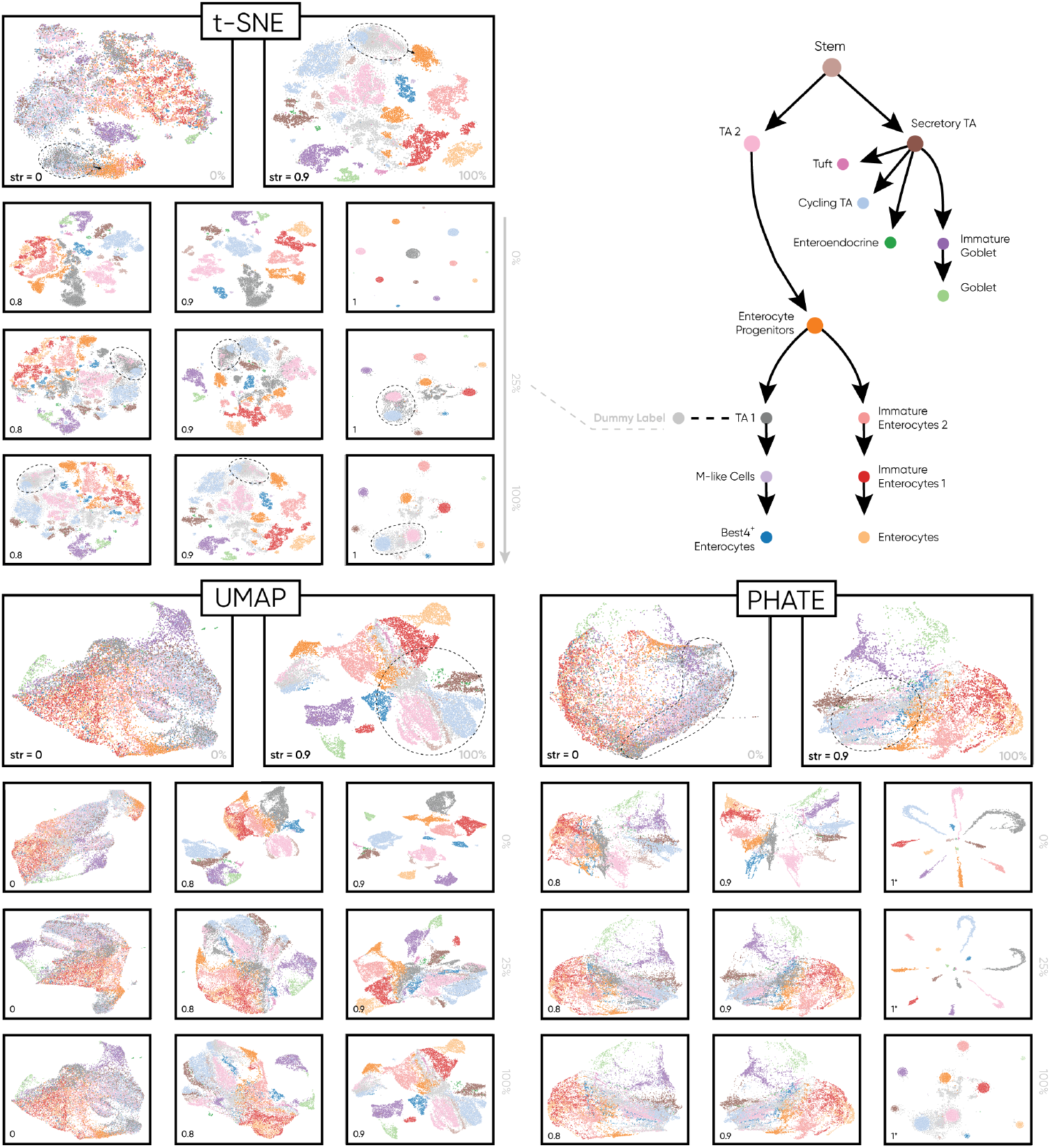
Haisu does not compromise the embedding of cells that do not have a label in the input graph. We depict 25% and 100% replacement of the TA-1 label with a ‘dummy’ label that is not present in the hierarchy across t-SNE, UMAP, and PHATE. Even at high strength values (*str*) of the hierarchical distancing factor, Haisu maintains relationships in the embedding circled in the figure. Notably, TA 1 cells remaining close to Cycling TA cells across the embeddings at 25% and 100% reduction despite their distance in the hierarchy graph. Thus, we do not comprise the integrity of each NLDR method, allowing for the observation of unknown classes in the context of a strongly influential, known hierarchy.

Across each method, the impact of the hierarchical distancing factor at different strengths is readily apparent in the progression of randomized embeddings. We tend to favor a lower strength factor to “push” clusters away from distant nodes in the hierarchy while maintaining the nuances of the raw NLDR methods. In this regard, the influence of information not present in the hierarchy is still expressed in the higher dimensional manifold and visible within the embedding. Thus, the input graph and strength factor encourage a user-centric approach that allows the input of prior knowledge. In this manner, one can manually adjust the strength factor to set the degree of separation when targeting specific relationships in the graph.

This approach can similarly be performed on each dataset for a gene-based hierarchy where sample classes are inferred with a given likelihood. Haisu in this mode can implement a probability-dependent approach that weights poor classifications less heavily. We first replace the ground-truth label of a sample *ℓ_k_* with an inferred label *ℓ′_k_* based on a given predictor. For example, we can predict by a cell’s expression of key genes per labels. Hierarchy nodes in this case could also be derived from representative proteins of the provided classes. Relevant UMI counts would first be summed for each node *υ* ∈ *V* of the gene-based graph following by the summation of key genes *g* ∈ *υ*. This simple classifier would finally label the given sample based on the node with the highest normalized expression.

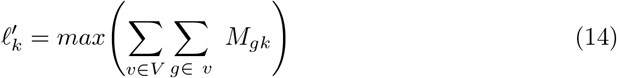

Scores for each inferred class can then be normalized and influence the likelihood-based pairwise distance matrix described in the Methods section. Because the graph is also weighted with a strength factor, we can control this effect by modulating str. Inferred labels with low UMI counts contribute less to the hierarchical pairwise distance matrix, ensuring that the original embedding is upheld. Thus, the original structure can be preserved when the confidence of a given prediction is low. This prevents faulty clustering from the influence of the hierarchical pairwise distance matrix. With 100% confidence for inferred labels, results of the gene-based method would mirror those of the class-based hierarchy (Fig. 1, 3).

## Discussion

Haisu formulates a direct relationship between the distance of two graph nodes in the hierarchy and the resulting pairwise distance in high-dimensional space (*θ · d* respectively). We chose this approach to ensure that closely related sibling nodes are encouraged in the embedding and farther nodes produce a noticeable amount of spreading. However, the effect of the hierarchy can easily be modified for a desired effect. To better express a hierarchical effect, one could introduce a diminishing factor such that nodes deeper into the graph have a reduced influence as opposed to the base classes. An application of such a strategy is relevant with the presence of distinct classes containing many nuanced subclasses as children. Another simple modification would be the application of a weighted graph in the calculation of the shortest path distance 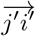 between labels *i*′ and *j*′.

The crux of Haisu’s practical use is in the choice of an appropriate strength factor. Primarily, one should employ this variable to control the extent of hierarchical distancing desired. We found its effect to be highly dependent on the characteristics of the dataset and the size/structure of the input hierarchy. In the included figures, we highlight lower strength factors that result in an embedding with similar visual structure to the base DR technique. Empirically, we suggest choosing a strength prior to significant clumping of neighboring nodes in the hierarchy graph. As described in the Methods section, our distancing algorithm is subtractive in nature. Rather than modifying the distance factor by an arbitrary amount, we perform a linear interpolation from 0 to 1 based on the relative graph node distances. We suggest that this strategy is more effective in maintaining the local and global structure of the data at all choices of strength factor.

We chose to modify the pairwise distance function prior to embedding due to the ease of generality into multiple NLDR methods. Additionally, due to possible random initialization that follows the initial pairwise distance calculation in most methods, label ordering can quickly become impossible to assign. However, with further modification of the base NLDR algorithms, one could instead modify the distance function, influence the initial random embedding, or extend the objective function with a hierarchical factor. We also chose not to integrate directed graphs with this approach to support more complicated inter-class relations, but one could extend the approach to punish a pairwise-comparison when moving against the directed graph. This would result in more inter-class separation and less intra-class separation which could be valuable for some datasets.

Though not investigated in this study, we suggest that for datasets in domains outside of scRNA-sequencing where labels assignments can be given with 100% certainty, Haisu would be effective even at higher dimensions for clustering. We suggest that clustering and processing at higher dimensions is feasible and could be an effective, supervised intermediate step because Haisu preserves the dimensionality reduction-specific architecture.

We proposed a novel general-purpose approach for nonlinear dimensionality reduction that incorporates an input hierarchy. We modify the premier visualization-centric techniques in this field and demonstrate our results on real-world single-cell datasets. Furthermore, we introduce a feature-based modification that enables users to integrate our method with weighted class labels.

## Data Availability

The authors declare that the data supporting the findings of this study are available within the paper and its supplementary information files.

## Code Availability

https://github.com/Cobanoglu-Lab/Haisu

## Acknowledgments

We thank Didem Çobanoğlu for her extensive guidance in the formulation of this approach and relevant biological expertise. We also thank Venkat Malladi and Spencer Barnes for assistance in selecting the single-cell RNA sequencing datasets and visualization methods.

---

**Figure S1.**
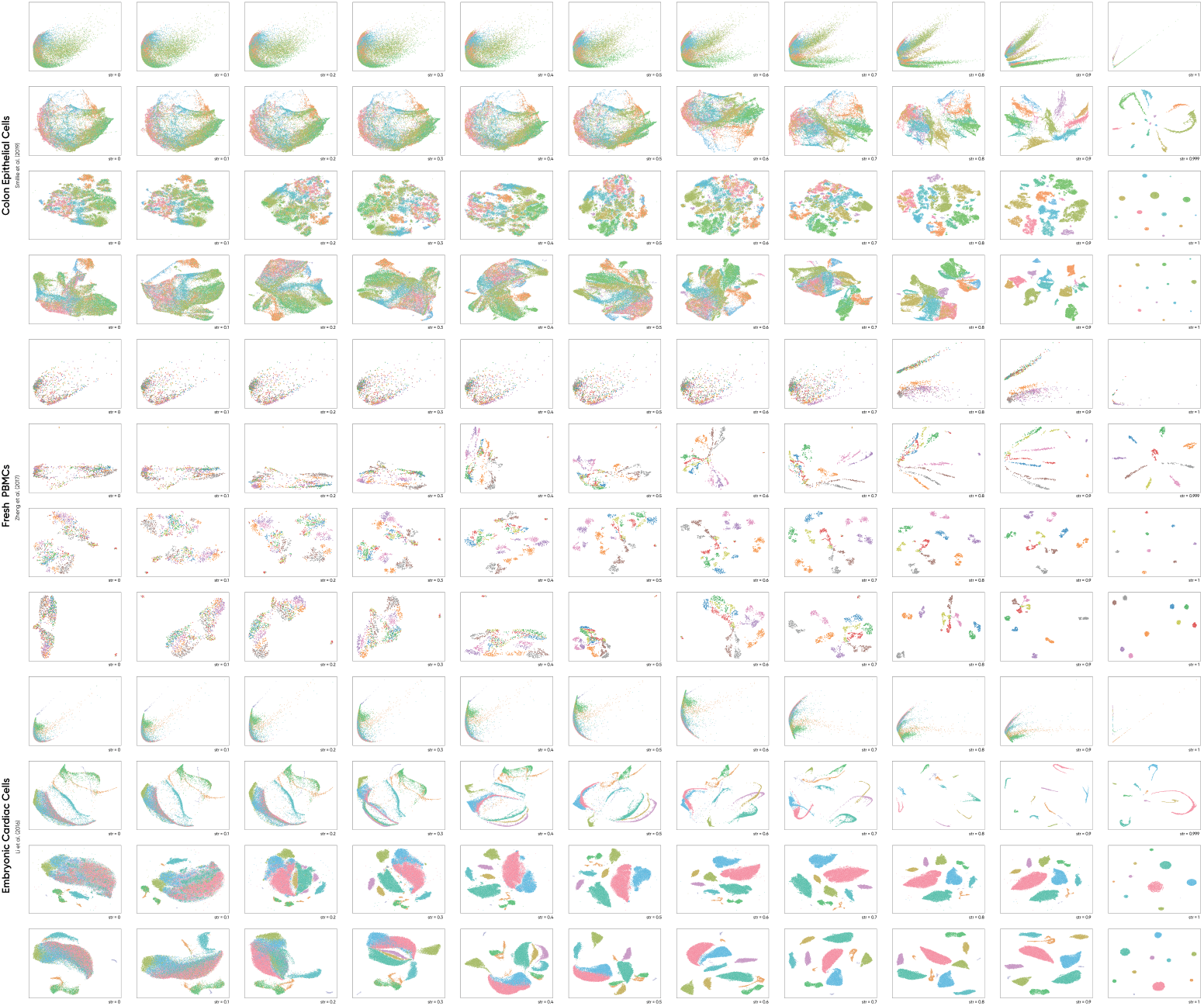
We display the full progression of Haisu from a strength factor of *str* = 0 to *str* = 1 corresponding to Fig. 1, 2, 3. Modified PCA, PHATE, t-SNE, and UMAP are displayed from top to bottom for each dataset.

**Figure S2.**
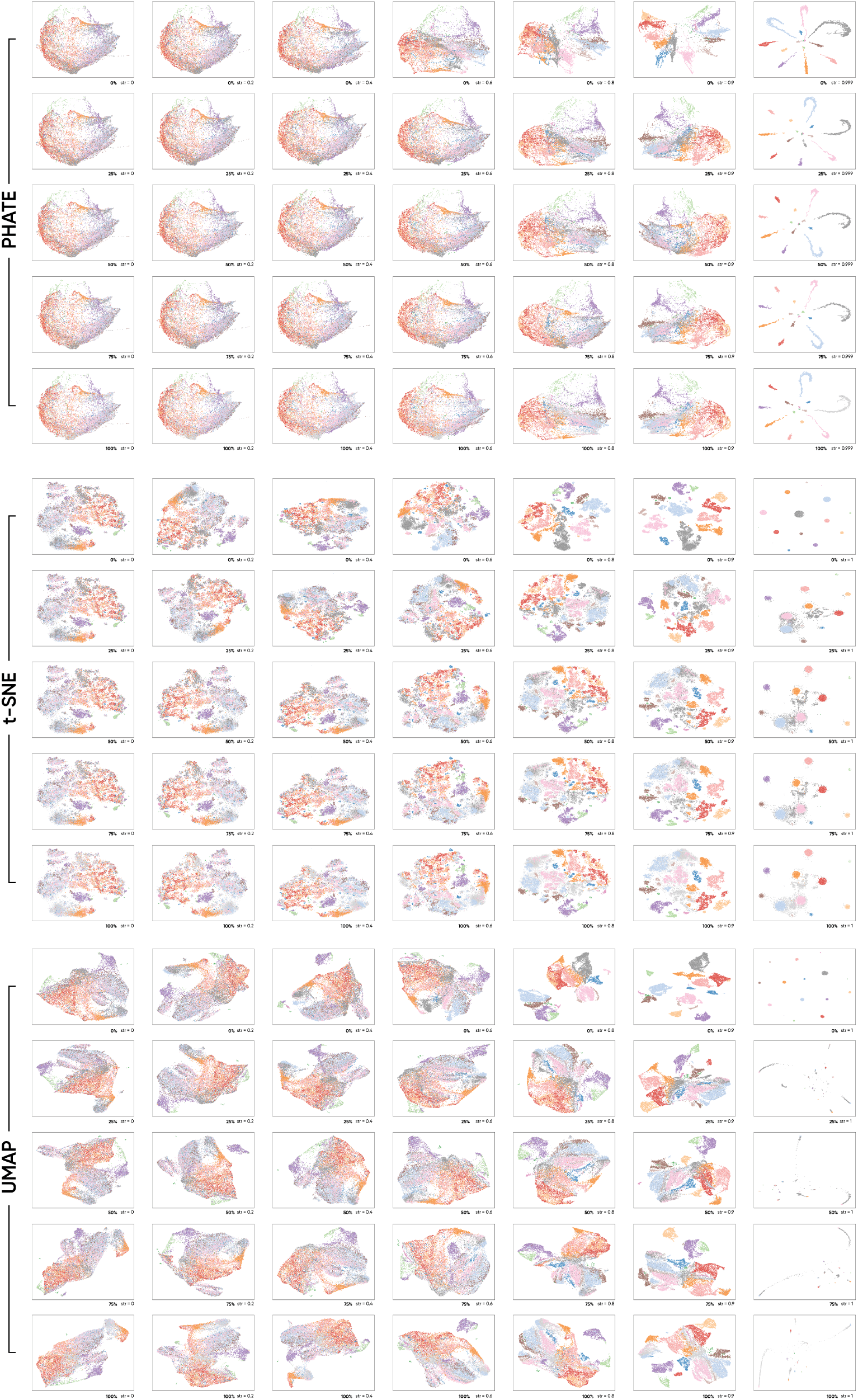
Full progression of Haisu strength factor values and percentage removal of the TA-1 label corresponding to Fig. 4. Modified PHATE, t-SNE, and UMAP nonlinear dimensionality reduction techniques are depicted.

